# Synchronization dependent on spatial structures of a mesoscopic whole-brain network

**DOI:** 10.1101/319830

**Authors:** Hannah Choi, Stefan Mihalas

## Abstract

We study how the spatial structure of connectivity shapes synchronization in a system of coupled phase oscillators on a mammalian whole-brain network at the mesoscopic level. Complex structural connectivity of the mammalian brain is believed to underlie the versatility of neural computations. The Allen Mouse Brain Connectivity Atlas constructed from viral tracing experiments together with a new mapping algorithm reveals that the connectivity has a significant spatial dependence: the connection strength decreases with distance between the regions, following a power law. However, there are a number of residuals above the power-law fit, predominantly for long-range connections. We show how these strong connections between distal brain regions promote rapid transitions between highly localized synchronization and more global synchronization as the amount of dispersion in the frequency distribution changes. This may explain the brain’s ability to switch rapidly between global and modularized computations.

## I. INTRODUCTION

Structural neural connectivity and its implications on the brain function have been a long-sought subject in neuroscience. Many previous studies have been limited either to small networks of few cells or coarser connectivity among larger brain regions [1–9], often binarized and without spatial information. Recent development of the Allen Mouse Brain Connectivity Atlas from anterograde florescent viral tracing experiments [10] provides us the unique opportunity to investigate precise weighted anatomical connectivity of the mammalian whole brain network. Combining the mesoscopic connectivity data with spatial information of the network, we seek a parsimonious representation of the weighted whole-brain network that captures salient network properties. Specifically, we investigate whether the network can be compactly represented solely based on the spatial dependence of the network topology.

Biological networks are inherently spatially-constrained. Recent studies have shown that geographic constraints play a critical role in generating graph properties of real-world neuronal networks [5, 11–20], which cannot be fully captured by classical generative network models such as the small-world network [2] and the scale-free network [21]. Yet, many of the studies are limited to binarized networks [11, 12, 17, 19, 20], and are focused explicitly on comparing graph theoretical measures [11, 13–20]. In this paper, we examine spatial embedding of the weighted whole-brain connectivity, and ask whether spatial dependence alone can depict the full computational capability of the brain network by studying dynamics of the network.

By analyzing the latest connectivity data from a new mapping algorithm, we find that the network connectivity strongly depends on its spatial embedding, with spatially close brain regions strongly connected and distal regions weakly connected. We study the precise relationship between connectivity and distance, and investigate possible computational roles of positive residual connection strengths that are not captured by the spatial dependence. To probe possible implications of the residual connections on the network dynamics, we construct a network of phase oscillators with the data-driven adjacency matrix, and compare its dynamics to those of the oscillator network with the connections strictly dependent on distance. We analyze spatial structures of synchronization by measuring the order parameter for varied amounts of dispersion in intrinsic oscillator frequencies. We further examine the strong connections between distal brain regions, by studying network dynamics when fractions of the strong residual connections are added to the spatially constrained network. Finally, we relocate the positive residuals to connections between nearby brain regions, increasing the connection strengths for the spatially close brain regions while eliminating strong distal connections. The network restructured this way maintains overall connection strength of the brain network but has the network structure different from that of the brain network. By comparing dynamics of such restructured network and the data-driven whole brain network, we show that the spatial locations of the strong positive residuals are important.

## II. RESULTS

### A. Spatial dependence of the mouse whole-brain connectivity

The mesoscopic mouse whole-brain connectivity was constructed based on viral tracing experiments available on Allen Mouse Brain Connectivity Atlas [10], with a recently developed interpolative mapping algorithm [22]. This produced a weighted and directed structural connectivity matrix with 244 brain regions as nodes. The data-driven mouse brain network is shown in Fig 1A, left column.

**Fig. 1.**
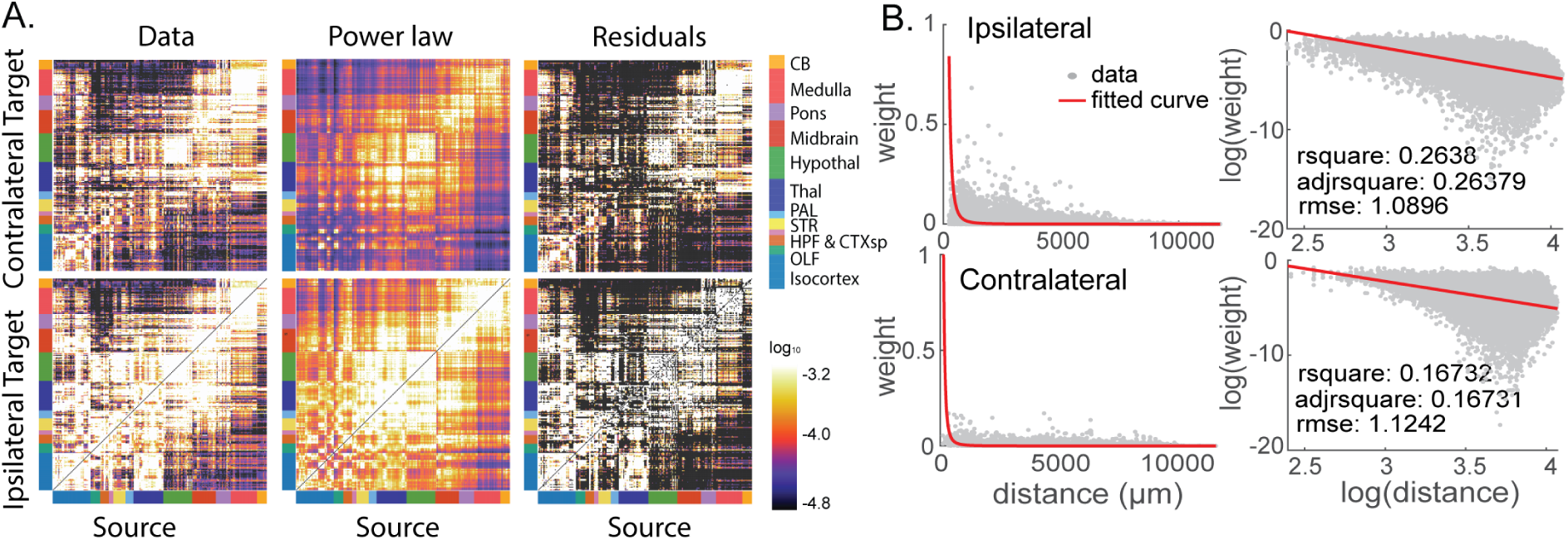
Connectivity matrices constructed from viral injection data and the power-law dependence on distance. (A) Connectivity matrix from viral tracing data (left); Reconstructed connectivity from the power-law dependence on distance between nodes (middle); residual connection strengths of the data-driven network above the power-law distance dependence. We show 244 brain regions divided in to coarser major brain divisions defined in the 3-D Allen Mouse Brain Reference Atlas. These divisions are: Isocortex, Olfactory Bulb, Hippocampus, Cortical Subplate, Striatum, Pallidum, Thalamus, Hypothalamus, Midbrain, Pons, Medulla, and Cerebellum. (B) Connection strengths as a function of distance between brain regions. The connections obtained from experiments (gray) are fit by a power law (red) on the log scale with base 10 (right panel). Inset: Goodness of fit.

We analyzed the relationship between the connection strength and spatial distance between brain regions in the data set. In accordance with previous studies on brain networks [5, 15–20], the connectome strongly depends on the spatial embedding; connections are stronger between spatially close regions and weaker between distal regions. Specifically, the connection strengths decrease with distances between brain regions following a power law (Fig 1B) rather than an exponential relationship, in agreement with previous studies on Allen Mouse Brain Connectivity data [18, 22].

We constructed adjacency matrices for the ipsilateral and the contralateral networks based on the power-law relationship, as shown in Fig 1A, middle column. While the general trend of decrease in connection strength with distance is clear and well-predicted by a power law, there are also a number of residual connection strengths that are not captured by the power-law relationship (Fig 1A, right column).

To understand the structure and effects of the residual connection weights that are not captured by the power-law dependence on distance, we had a closer look at these residuals. For both ipsilateral and contralateral connections, a long, positive tail is observed in the distribution of residual connections weights, suggesting strong distal connections above the power-law dependence on distance (Fig 2A, B). The strongest 20 residual connections are plotted in Fig 2C. We observed that for ipsilateral network, connections from preparasubthalamic nucleus (PST) to subthalamic nucleus (STN), laterodorsal tegmental nucleus (LDT) to Barrington’s nucleus (B), dorsal motor nucleus of the vagus nerve (DMX) to gracile nucleus (GR), cuneate nucleus(CU) to gracile nucleus (GR), and locus ceruleus (LC) to Barrington’s nucleus (B) are few examples of the strong distal connections unexplained by the power-law dependence on distance. For contralateral connectivity, on the other hand, many of the strongest residuals above the power-law relationship include the connections between the same regions in different hemispheres as well as connections to and from hippocampal areas.

**Fig. 2.**
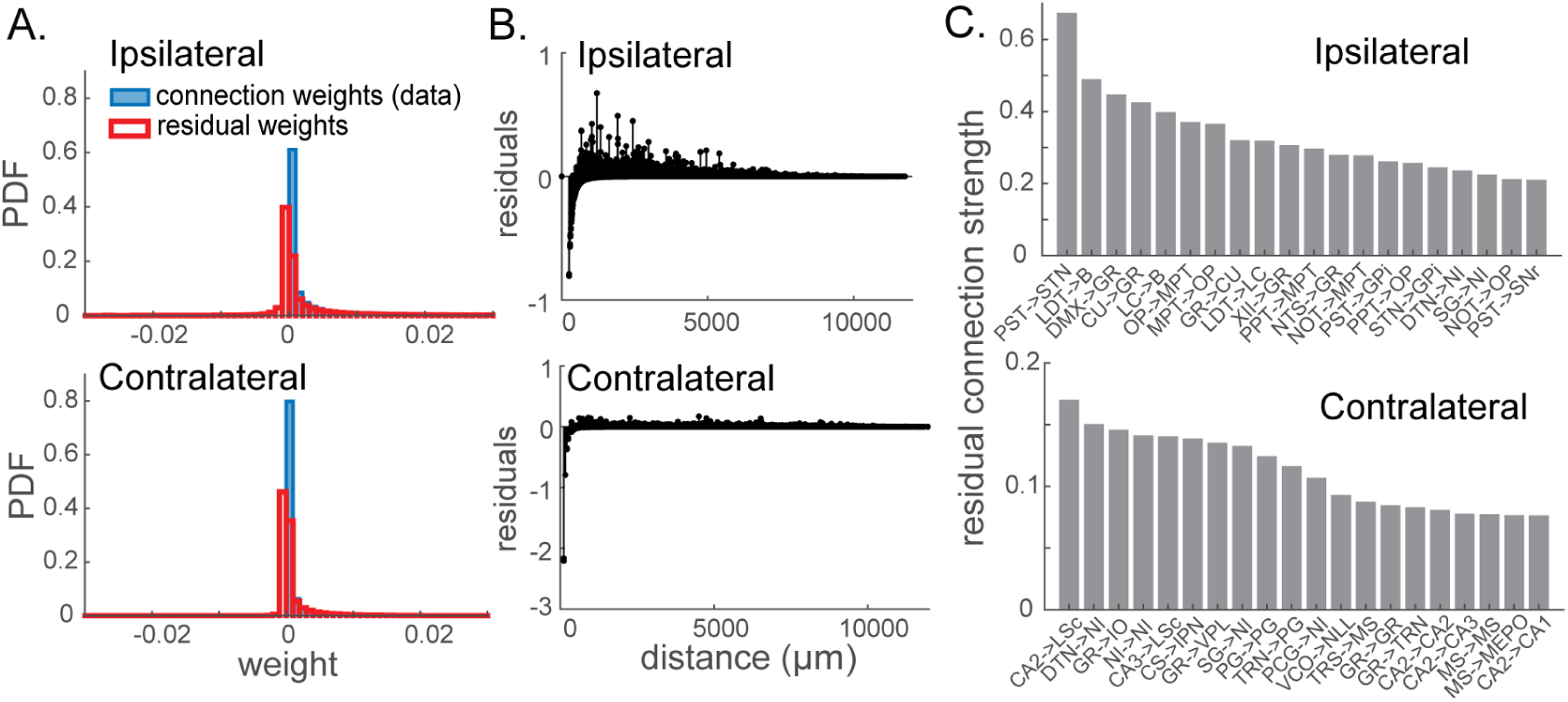
Residual connection weights unexplained by the power-law distance dependence. (A) Distributions of the connection strengths from the data (blue) and the residual connection strengths (red). (B) Residual connection weights as a function of distance between nodes. (C) Directed pairs of brain regions with large positive residual connections. These represent strongly-connected distal regions. For reference on the acronyms of the regions, see the Allen Mouse Brain Reference Atlas.

### B. Phase oscillators and network coherence measures

Do these positive residual connections between distal regions have any computational significance? In other words, can we capture the full computational capacity of the mesoscopic brain network with connectivity governed by strictly distance-dependent rules, with the residuals removed? To test this, we compare dynamics of the data-driven brain network to those of an artificial, strictly distance-dependent network generated by the power-law relationship. Specifically, we built a network of coupled phase oscillators whose coupling strengths are described by the weighted adjacency matrix of the data-driven brain network or the power-law distance-dependent connectivity. Each of these Kuramoto-type phase oscillators corresponds to a brain area. The phase of region *i*, represented by *θ_i_*, is described by:

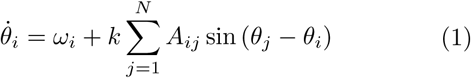

where *ω_i_* denotes the natural frequency, and *k* describes the coupling coefficient. *A_ij_* is the adjacency matrix of the network. For the case of the data-driven brain network, 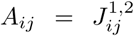 where 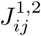 indicates the adjacency matrix obtained from viral tracing data, for the ipsilateral-only 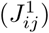 or both ipsilateral and contralateral 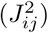 connections. For simulations of the artificial, distance-dependent network, 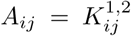 indicates the adjacency matrix constructed by making the connection weights strictly follow the power-law dependence on distance, for the ipsilateral-only 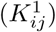 or both ipsilateral and contralateral 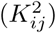 connections. *N* denotes the number of nodes of the network. We use either *N* = 244 or *N* = 488, for simulations of one hemisphere with ipsi-lateral connections only and simulations of both hemispheres with ipsilateral and contralateral connections, respectively. The natural frequencies *ω_i_* were randomly chosen from a symmetric, unimodal distribution *g*(*ω*). In this paper, we used a Gaussian distribution with the mean at 0 and the standard deviation *σ* for *g*(*ω*), as done in other studies [23–26]. Note that we can use zero mean without loss of generality as the mean frequency can be shifted by introduction of the change of variables. We also tried an uniform distribution with the zero mean bounded by *σ*, which yielded the same results as with Gaussian distributions.

We investigated the dynamics of the data-driven network and the power-law generated network using Eq. 1, and measured the network coherence by calculating the “universal” order parameter, recently proposed in [26]. Unlike the original order parameter proposed by Kuramoto [27, 28] (see Eq. 2 in Methods), the universal order parameter [26] can be used to quantify coherence in more general, weighted networks of oscillators, without assuming all-to-all connectivity. The universal order parameter accounts for the network topology and its influence on the phase coherence. Therefore, we can compare network coherence in topologically different weighted networks even when their total connections strengths are not the same. Furthermore, the universal order parameter captures partially phase-locked states accurately. To quantify different degrees of network coherence and to visualize localized and global synchrony, we measured the universal order parameter both for the whole network of oscillators (Eq. 3 in Methods) as well as for subnetworks of different spatial scales (Eq. 4 in Methods). By computing the order parameter for the subnetworks, we describe the order parameter as a function of distance.

Obtaining an explicit, analytical relationship between the order parameter and generalized network structures has been a challenging problem in studies of phase oscillators on complex networks [29, 30]. While analytical expressions for the order parameter as a function of the adjacency matrix have been derived in previous works, these mean-field approaches are based on strong assumptions of a large network with sufficiently high average degree, valid only near the onset of synchronization [24, 25, 27, 31, 32]. Existing analytical approaches, therefore, are not applicable to the complex mesoscopic brain network of a finite size. We thus address the relationship between the network coherence and the network structure by computing the order parameter based on numerically obtained time series of the oscillators.

The initial conditions for Eq. 1 were set to zero frequencies, and Eq. 1 was integrated numerically using 4th order Runge-Kutta method, with discrete time step ∆*t* =1 for *N_t_* = 10^3^ steps, until a stationary state is reached. We confirmed that a stationary state is obtained with *N_t_* steps, by testing larger (up to *N_t_* = 10^4^) numbers of time steps which did not alter the results. The data from the first *N_t_* /2 steps are discarded in measuring the order parameter. We computed the order parameter using Eq. 4 in Methods for varied amounts of perturbation represented by standard deviation *σ* of the natural frequency distribution, with fixed coupling coefficient *k* = 0.2. Note that increasing the coupling coefficient *k* with fixed *σ* has qualitatively the same effect as decreasing *σ* with *k* fixed, as the ratio of *k/σ* determines the network coherence. For each amount of dispersion *σ*, we performed 5 independent runs with different configurations of the intrinsic frequencies randomly sampled from the probability distribution *g*(*ω*), and plotted the average and the standard deviation of the order parameter, as a function of distance between nodes (Fig 3B) as well as a function of dispersion amount in the intrinsic frequencies *σ* (Fig 3C).

**Fig. 3.**
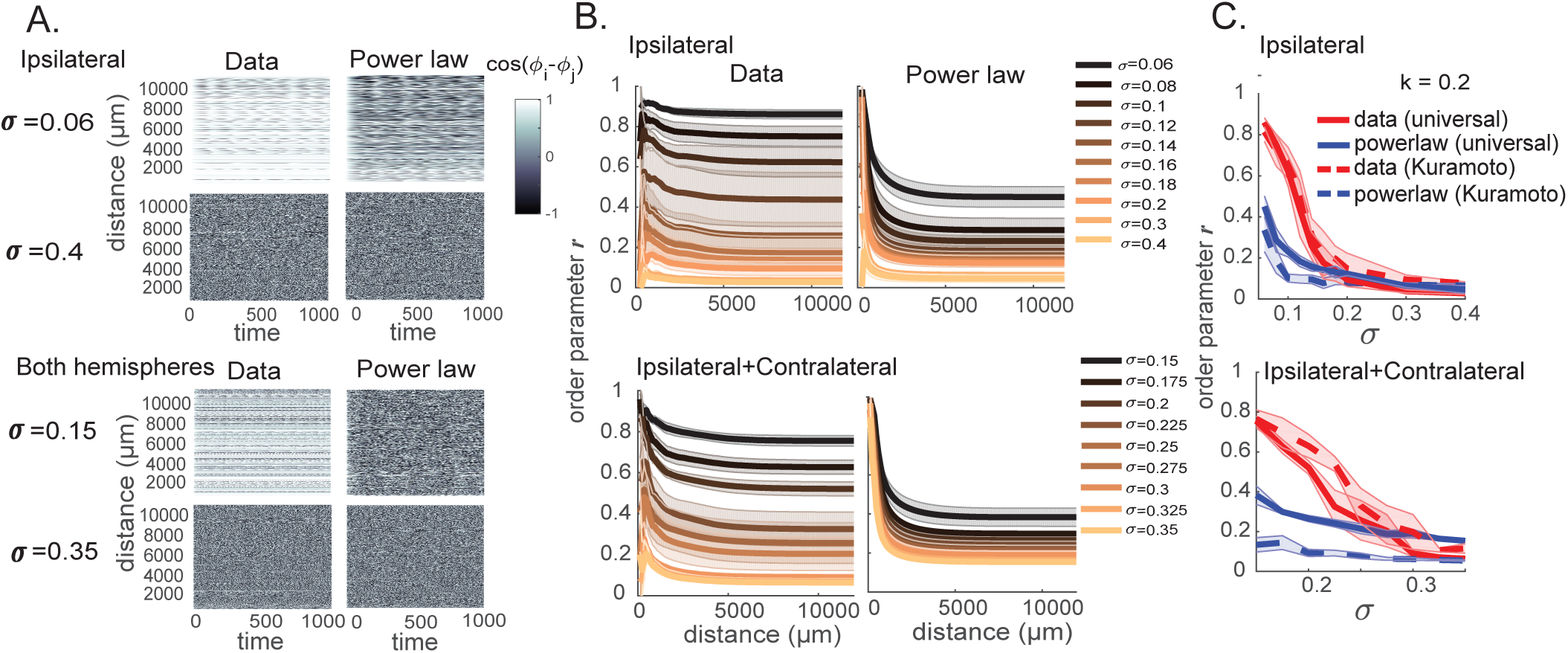
Local and global synchronization of the data-driven brain network and the power-law network. (A) Phase differences cos(*θ_j_* − *θ_i_*) between pairs of nodes (*i,j*) as a function of time (x-axis) and distance between the pairs (y-axis) for the data-driven and power-law networks. For both networks, the same set of natural frequencies generated from normal distributions with zero mean and standard deviation *σ* was used. (B) Universal order parameter *r* for subnetworks at different spatial scales, for the data-driven and the artificial power-law networks under different amounts of perturbation. Order parameter *r* is averaged over 5 generations of sets of natural frequencies. (C) Universal order parameter (solid) and Kuramoto’s original order parameter (dotted) for the whole networks of the data-driven connectivity (red) and the power-law distance-dependent connectivity (blue), as a function of perturbation amount *σ*. Coupling coefficient is fixed at *k* = 0.2. Lines and shades correspond to the mean and the standard deviation over multiple generations of intrinsic frequencies, respectively.

### C. Sensitivity of the network coherence

In Fig 3A, we show the phase difference cos(*θ_j_* − *θ_i_*) between pairs of nodes (*i,j*) plotted against time and distance between the pairs. Interestingly, for the same amount of change in perturbation ∆*σ*, the real brain network switches between an asynchronous state and global synchrony, while the power-law governed network fails to make such a drastic change in synchronization state. This difference is manifested in the order parameter. Fig 3B shows the universal order parameter (Eq. 4) for subnetworks of different spatial ranges. In the data-driven brain network, increasing the frequency dispersion a results in a transition from global synchrony to localized coherence (Fig 3B, left column, Data). However, in the artificially generated, strictly distance-dependent network, the same amount of perturbation change does not induce such a leap in the network coherence state as in the real brain network (Fig 3B, right column, Power law).

Such trend can be also visualized in the order parameter for the whole network. The overall universal order parameter increases with decreasing frequency dispersion in both the data-driven and the power-law networks (Fig 3C). However, the change in order parameter is significantly larger in the data-driven brain network. This trend appears in both the single hemisphere network with only ipsilateral connections and the whole brain network with both ipsilateral and contralateral connections. For comparison, Kuramoto’s original order parameter (Fig 3C, dotted) was also plotted. Because the original Kuramoto’s order parameter assumes the same all-to-all connectivity and does not measure coherence scaled to the overall degree of the network, we see that the Kuramoto order parameter is lower than the universal order parameter for the power-law network. Nevertheless, for either type of order parameter, we observe that the data-driven brain network spans a larger range of coherence states than the power-law governed network. This indicates that in the real brain network, a small perturbation in intrinsic frequencies induces a rapid transition between highly localized network synchrony and a more globally synchronized state, while in the network with connections strictly following a power-law dependence on distance, such a rapid transition between local and global coherences is not possible. Therefore, the residual connection strengths that are not explained by the spatial dependence rule may have some computational significance, enabling even small perturbations to induce a switch between global and modularized computations.

### D. Effects of strong long-range connections

We next examined what aspects of the residual connection strengths confer the network ability to span a wide range of coherence states. In previous studies on coupled oscillators, it has been found that even a small fraction of shortcuts in a small-world network significantly improves synchronization of the network [23, 29]. Motivated by this, we hypothesized that positive residual connections, namely, strong connections between distal brain regions, underlie the rapid transition between localized and global network synchronies. We tested this hypothesis by re-introducing small fractions of the positive residuals to the power-law distance-dependent network. As manifested in Fig 4, adding just a small fraction (top 5 percentile) of the strongest positive residuals to the power-law generated network recovers the steep decrease in order parameter with growing perturbation (Fig 4, cyan). As the fraction of positive residuals included in addition to the power-law network increases, the trend in the order parameter resembles more of that of the real brain network (Fig 4, red).

**Fig. 4.**
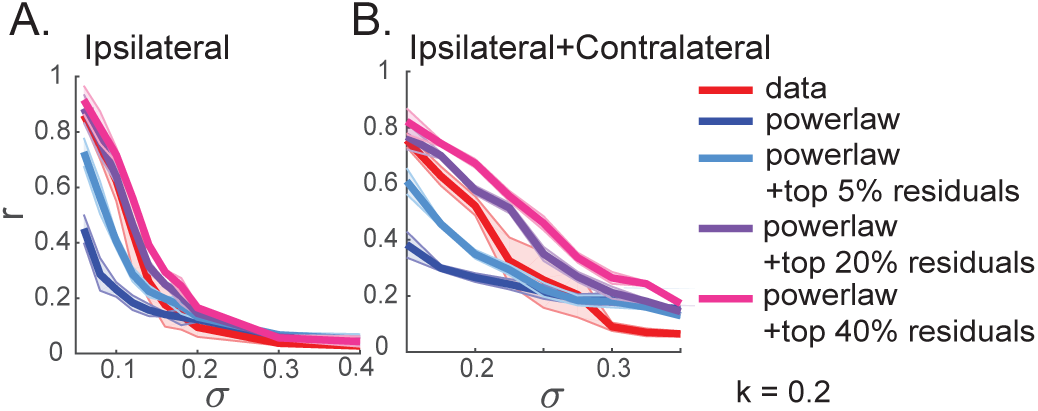
Order parameter *r* for the power-law distance-dependent network with a fraction of the residual connections added. Whole network order parameter *r* as a function of perturbation *σ*, for networks constructed by adding different percentiles of positive residual connections to the power-law network of (A) ipsilateral and (B) ipsilat-eral+contralateral connections.

Does the location of these strong connections have any significance in emergence of the rapid phase transition? To test whether the sensitivity of the network coherence to frequency dispersion can be recovered by simply adding the positive residuals anywhere to increase the overall connection strength of the power-law network, we studied the dynamics of the network constructed by relocating the positive residuals. We generated two networks with positive residuals relocated. In one of them, the positive residuals above the power-law relationship were positioned at random locations on the network (shuffled). In the other, the positive residual connections were relocated and added to connections between spatially close regions, by distributing the total positive residual connection strength among the connections between nodes within 500*μm*. The resulting networks thus maintain the total connection strengths of the real brain network, but have altered network structures. When the locations of the positive residuals are shuffled and thus there are strong connection weights between distal brain regions, the dependence of network synchronization on *σ* remains similar to that of the data-driven network, as portrayed by the order parameter in Fig 5 A,C, in gray and Fig 5 B,D, left. In other words, although the precise network structure is different from that of the data-driven net-work, the network with shuffled residuals maintains its sensitivity to perturbation, rapidly changing between localized and globally coherent states. However, when the positive residuals are relocated to proximal connections, the network coherence is no longer sensitive to small perturbations in the frequency distribution (Fig 5 A,C, green; B,D, right), in spite of the unaltered total connection strengths. Unlike the network with randomly relocated residuals, the network with positive residuals relocated to proximal connections lack strong connections between distal brain regions.

**Fig. 5.**
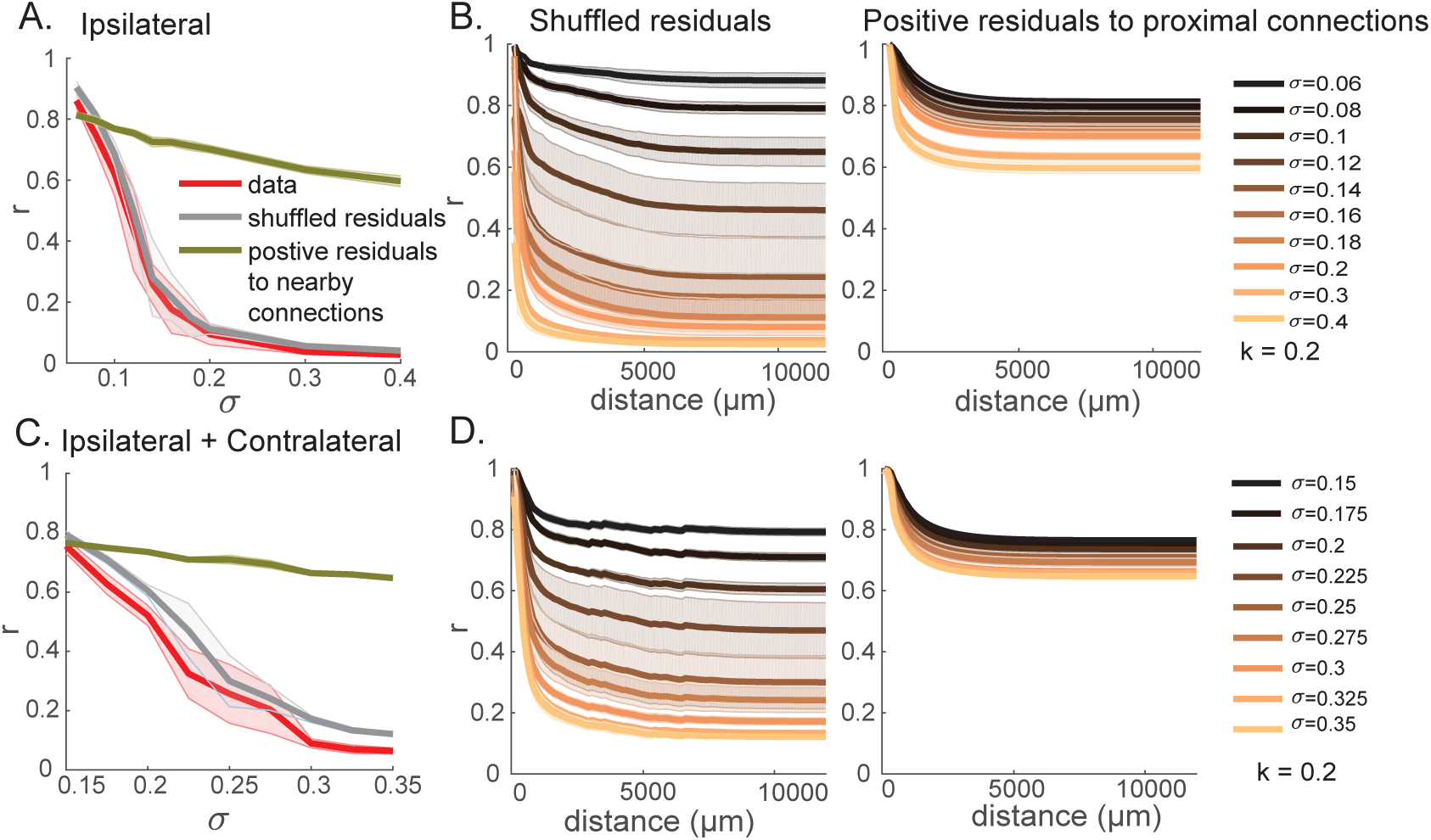
Synchronization measured when the network structure is altered but the overall degree remains the same as the data-driven network. (A,C) Whole network order parameter *r* as a function of perturbation amount *σ*, for the network generated by adding the residual connection weights to random locations (gray) and the network constructed by relocating positive residuals (averaged) to connections between spatially close regions (< 500*μm*) (green). The order parameter for the data-driven brain network (red) is shown for comparison. (B,D) Order parameter as a function of spatial scale of subnetworks, for the networks constructed by shuffling locations of residual connections (left) and by relocating positive residuals to nearby connections (right). Measures for a single hemisphere with ipsilateral connections (A, B) and both hemispheres with ipsilateral and contralateral connections (C, D) are shown.

This shows that the location of strong connections above the power-law dependence on distance is critical for generating a steep change in order parameter; however, the precise positions of the strong connections do not have to match those of the data-driven network, as long as there is a sufficient amount of strong connections between distal nodes. In sum, the spatial structure of the network connectivity plays a key role in maintaining the brain’s ability to change its computational states with small perturbations, and such sensitivity cannot be achieved by simply matching the total network connection strengths. However, the structure does not have to precisely match that of the real brain network. What is critical to maintain, rather, is some strong connections between distal regions.

## III. CONCLUSION

In this paper, we studied synchronization of a spatially constrained model of a weighted whole-brain network at mesoscale, constructed from viral tracing experiments. We found that the connectivity has a significant spatial dependence, with the connection strength decreasing with distance between the regions following a power law. However, by studying the network dynamics of phase oscillators, we found that a network gener

ated by the simple spatial constraints alone cannot reproduce the full computational versatility of the mesoscopic whole-brain network. Rather, we need to consider additional complexities of the network structure to capture their possibly significant roles in neural computation. Specifically, we found that residual connections not explained by the power-law dependence on distance have a long positive tail, corresponding to strong connections between distal brain regions. By computing the recently proposed universal order parameter, we showed that these strong distal connections underlie sensitive dependence of network synchrony on perturbation, potentially responsible for the brain’s exceptional ability to change its computational states depending on stimulus and behavioral context. Furthermore, our analyses on a network constructed by adding a small fraction of strong positive residuals to the spatially-constrained connectivity, as well as a network with the positive residuals relocated to random locations and proximal connections, reveal the key element underlying the rapid switch between global and modularized synchronies - strong connections between distal brain regions. In other words, the network’s sensitivity to perturbation cannot be reproduced by simply manipulating the overall connection strengths, as locations of positive residual connections should be taken into consideration. A spatially-constrained model plus an idiosyncratic sparse matrix which features strong connections between distal regions, provides a parsimonious representation of the measured connectivity.

We hypothesize that the sharp transition between locally and globally synchronized states in the data-driven network, which is absent in the spatially-constrained power-law model, may underlie the brain’s ability to rapidly switch between global and modularized computations [33]. Such feature is known to be impaired in the brain under pathological conditions such as Alzheimer’s disease, suggested by studies showing more modular structures and decreased global efficiency in brain connectivity constructed from EEG, MEG, fMRI, and diffusion tensor tractography [34–37]. Moreover, there is an experimental evidence for disruption of long-range connections in Alzheimer brain network [36], in agreement with our model results. A more detailed future study on genetically-controlled mouse models of Alzheimer’s disease will shed light on the possible link between changes in structural connectivity and impairment in rapid phase transitions of the whole-brain network.

In this paper, we infer the dynamics of the mesoscopic brain network by constructing a network of phase oscillators with the coupling strengths determined by the structural connectivity obtained by viral tracing experiments. Thus, while the structural connectivity is based on actual data, the dynamics we conferred on the network are arbitrary. Building a more realistic, data-driven dynamic network based on imaging experiments such as calcium-imaging or ECOG will be a crucial future extension of our study of connecting the network structures to the network dynamics. However, our simulations with phase oscillators, despite their generality, still make valuable predictions on computational roles of spatial structures of the mesoscopic whole-brain network, underlining the importance of distal connections on the network dynamics.

## IV. METHODS

### A. Mouse whole-brain connectivity data

The mesoscopic mouse whole-brain connectivity was obtained from Allen Mouse Brain Connectivity Atlas (http://connectivity.brain-map.org/), constructed based on anterograde viral tracing experiments in wild type C7BL/6 mice [10]. Based on the experimental data, a recently developed interpolative mapping algorithm was used to construct a model of whole brain connectivity at the 100 *μm*-voxel scale [22]. The voxel-based connection strengths were averaged over each brain region to produce a connectivity matrix with 244 brain regions per hemisphere as nodes, larger than the adjacency matrix of 213 pairs of nodes previously obtained from the linear model in [10]. For elements of the connectivity matrix, we use the *normalized projection density*, defined as the connection strength between two regions divided by the volume of the source and target regions. In order to account for the size of the source region, we also studied the relationship between the connection strength divided only by the size of the target region and the distance between two regions. In this case, however, the fit to either a power law or an exponential function was not very good which is not surprising given that the connection strength that are not fully normalized with respect to the size of the source and the target is not an intrinsic quantity. For more details on the viral tracing experiments and the interpolative algorithm used to construct the connectivity matrix, see [10] and [22]. The connectivity matrix was first normalized to have values between 0 and 1. For the ipsilateral connection matrix, the diagonal entries were set to 0 removing self-connectivity, as done in [4, 20].

### B. Dependence of connection strengths on interregional distance

We fitted connection strengths as a function of interregional distance, where the distance between each pair of nodes was determined by computing the Euclidean distance in 3-dimensional coordinates between the centroids of the brain regions. Specifically, power-law functions for relationships between connection strength and interregional distance were fitted by using least squares on the log scale. For each of the ipsilateral and contralateral connectivity matrices, we found *α* and *β* by fitting the data to 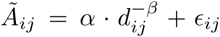, where *Ã_ij_* denotes the connection strength from node *j* to node *i, d_ij_* indicates the distance between nodes *i* and *j*, and *ε_ij_* is the residual error. We obtained *α* = 6.92 × 10^6^ and *β* = 2.886 for ipsilateral connectivity, and *α* = 6.71 × 10^5^ and *β* = 2.685 for contralateral connectivity (Fig 1B). In agreement with previous studies on Allen Mouse Brain Connectivity data [18, 22], we found that the power law explains the relationship slightly better than the exponential dependence (ipsilateral r-square: 0.264 vs 0.257, rmse: 1.089 vs 1.095; contralateral r-square: 0.167 vs 0.135, rmse: 1.124 vs 1.146).

We also investigated the power-law constrained network where the relationship between connection strength and interregional distance was found on the real scale, using nonlinear least squares (Levenberg-Marquardt algorithm), which has a poorer explanatory power than linear least squares on the log-scale (r-square: 0.264 vs 0.157 (ipsilateral) / 0.167 vs 0.131 (contralateral)). While this method generated a different power-law function from the one found by least squares on the log-log scale, the dynamics on the power-law network obtained by using nonlinear least squares maintained the same core characteristics, distinct from the data-driven brain network-the order parameter is less sensitive to the dispersion amount in intrinsic frequencies.

### C. Order parameter

In this section, we describe order parameters that were proposed previously [24, 25, 27, 28], demonstrating advantages of the recently developed universal order parameter [26] in our analysis.

In order to quantify a transition from incoherence to synchronization in the original model of phase oscillators with all-to-all connectivity, Kuramoto introduced an order parameter (*r*_Kuramoto_) that measures the average of phase differences of all pairs of oscillators [27, 28], such that:

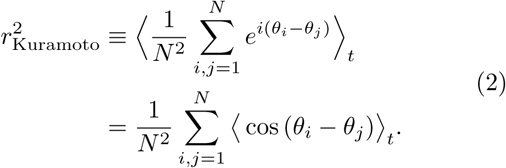

*< … >_t_* denotes the average over time. However, this unweighted order parameter is not a good measure when comparing collective synchronizations in two networks described by different connectivity matrices, as it does not capture the topology of the networks.

To extend the use of order parameter to more general, weighted networks of oscillators, Restrepo et al [24, 25] proposed an order parameter which is defined as the average of local order parameters which measure the coherence of the inputs to each node. This parameter, however, does not capture partially phase-locked states well. Recently, Schroeder et al [26] proposed a new, “universal order parameter” which overcomes the shortcomings of the previous order parameters. This newly proposed universal order parameter is defined as:

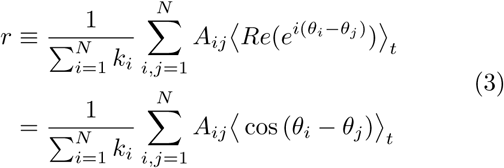

where 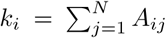 is the input strength of node *i*. Note that in unweighted binary networks, this measure represents in-degree [4]. This order parameter accounts for the network topology and its influence on the phase coherence, enabling a fair comparison between two topologically different weighted networks even when their total connection strengths are not matched. As this universal order parameter accurately captures partial synchrony within the network, different degrees of synchronization can be measured by order parameter of the whole network.

Furthermore, degree of coherence as a function of spatial extent can be obtained by computing the order parameter for subnetworks of different spatial scales. The order parameter *r* can be described as a function of distance *d*:

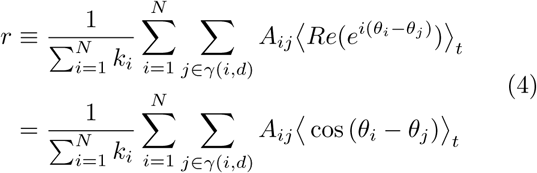

where *γ*(*i, d*) indicates the set of nodes within spatial distance *d* from node *i*. 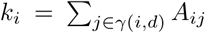 is the total connection strength of node *i* when the subnetwork composed of nodes within distance *d* from node *i* is considered. The order parameter of the whole network is obtained when *d* = size of the network (11752 *μm* for ipsilateral and 11955 *μm* for contralateral connectivity).

## ACKNOWLEDGEMENT

We thank Joseph Knox for providing the connectivity matrix used in this analysis. We also thank Kameron Harris for many helpful comments and suggestions. This work was supported by the Allen Institute for Brain Science. We wish to thank the Allen Institute founders, Paul G. Allen and Jody Allen, for their vision, encouragement, and support. Part of this work was done while H.C. was visiting the Simons Institute for the Theory of Computing.

